# What determines trophic niche breadth? A global analysis of freshwater fishes using isospaces

**DOI:** 10.1101/2025.05.27.656387

**Authors:** Friedrich Wolfgang Keppeler, Tommaso Giarrizzo, Carmen G. Montaña, Alphonse Adite, Şenol Akın, Ronaldo Angelini, Caroline C Arantes, Evanilde Benedito, Thethela Bokhutlo, Rana El-Sabaawi, Alexandre Garcia, Ivan Gonzalez-Bergonzoni, David Hoeinghaus, Joel Hoffman, Olaf P. Jensen, Erik Jeppesen, R. Keller Kopf, Craig A. Layman, Edwin Orlando Lopez, Bryan M. Maitland, Shin-ichiro S. Matsuzaki, Jill A. Olin, Gordon Paterson, Yasmin Quintana, Carlos Eduardo de Rezende, Ashley Trudeau, Paulo A. Trindade, Thomas F. Turner, Eugenia Zandona, Kirk O Winemiller

**Affiliations:** Federal University of Ceará, Brazil; Federal University of Pará, Brazil; Federal University of Rio Grande do Norte, Brazil; Stephen F. Austin State University, USA; Université d′Abomey-Calavi, Benin; Yozgat Bozok University, Türkiye; West Virginia University, USA; State University of Maringá, Brazil; Botswana International University of Science and Technology, Botswana; University of Victoria, Canada; Federal University of Rio Grande, Brazil; Universidad de la Republica, Uruguay; University of North Texas, USA; Environmental Protection Agency, USA; Center for Limnology, University of Wisconsin – Madison, USA; Aarhus University, Denmark; Charles Darwin University, Australia; Wake Forest University, USA; Industrial University of Santander, Colombia; USDA Forest Service, Rocky Mountain Research Station, USA; National Institute for Environmental Studies, Japan; Michigan Technological University, USA; Shedd Aquarium, USA; Northern Fluminense State University, Brazil; National Marine Fisheries Service, USA; University of New Mexico, USA; State University of Rio de Janeiro, Brazil; Texas A&M University, USA

**Keywords:** Climate conditions, Environment, Isotopic niche, Food webs, Functional traits, Macroecology, Stable isotope analysis, Trophic niche, Sampling scale

## Abstract

Trophic niche is one of the most tractable dimensions of a species niche. Several hypotheses have been proposed to explain global variation in species’ trophic niches, but empirical tests are limited. Stable isotope analysis (SIA) of δ¹³C (a proxy for basal sources) and δ¹ N (a proxy for vertical trophic position) has increasingly been used to estimate trophic niche variation. We performed SIA on a dataset of over 33,000 stable isotope samples from freshwater fishes across six ecoregions. For 541 populations (358 species), we estimated the size of δ¹³C × δ¹ N isospace—a bivariate representation of trophic niche breadth. We evaluated isospace variation in relation to environmental factors, species traits, and sampling scale, testing both novel and long-standing hypotheses about the drivers of trophic niche variation. A Boosted Regression Tree Model best predicted isospace, with environmental and trait-based variables among the strongest predictors. Fish isospaces were broader in regions with warmer temperatures, higher humidity, and precipitation variability, suggesting that more productive, diverse, and seasonal ecosystems are associated with broader trophic niches. Conversely, isospace size declined with increasing basin fish richness, which may reflect heightened interspecific competition or historical competitive exclusion. Isospace size was slightly smaller in large lakes compared to rivers, streams, and small lakes. Within-population variation in body size was strongly and positively associated with isospace size, highlighting the role of ontogenetic dietary shifts. Fish with truncate-rounded fins had broader isospaces than those with forked-lunate fins, likely due to the former having more limited movement and greater spatial isolation that resulted in greater between-individual niche variation. Primary consumers had larger isospaces than intermediate and top consumers.

Predators showed relationships consistent those observed from analysis encompassing all trophic guilds, but non-predators deviated with regards to four variables—solar radiation, basin richness, average body size, and caudal fin aspect ratio. Isospace size increased with the number of sampled individuals, habitats and years surveyed. Although isospace patterns have well-documented limitations as trophic ecology metrics, our findings nonetheless conform with several longstanding hypotheses for trophic niche variation and stimulate new ideas about environmental and biological drivers of niche breadth.

## INTRODUCTION

The niche concept is central to ecology and evolutionary biology (Baker et al., 2022) and underpins many theories in community ecology (e.g., Diamond, 1975; Leibold et al., 2004; Vellend, 2016). Trophic niche breadth describes the range of food resources used by an organism or population and is one of the most tractable and commonly studied aspect of the ecological niche (Bearhop et al., 2004). Population trophic niche breadth is shaped by both individual diet breadth, as well as dietary differences among individuals that may include trophic specialists and/or generalists (Bolnick et al., 2003; León et al., 2024). Several hypotheses have been proposed to explain variation in population trophic niches. Broadly, these hypotheses fall into three main categories: (i) environmentally driven, (ii) trait-driven, and (iii) scale-dependent (Table 1, Figure 1).

**TABLE 1.**
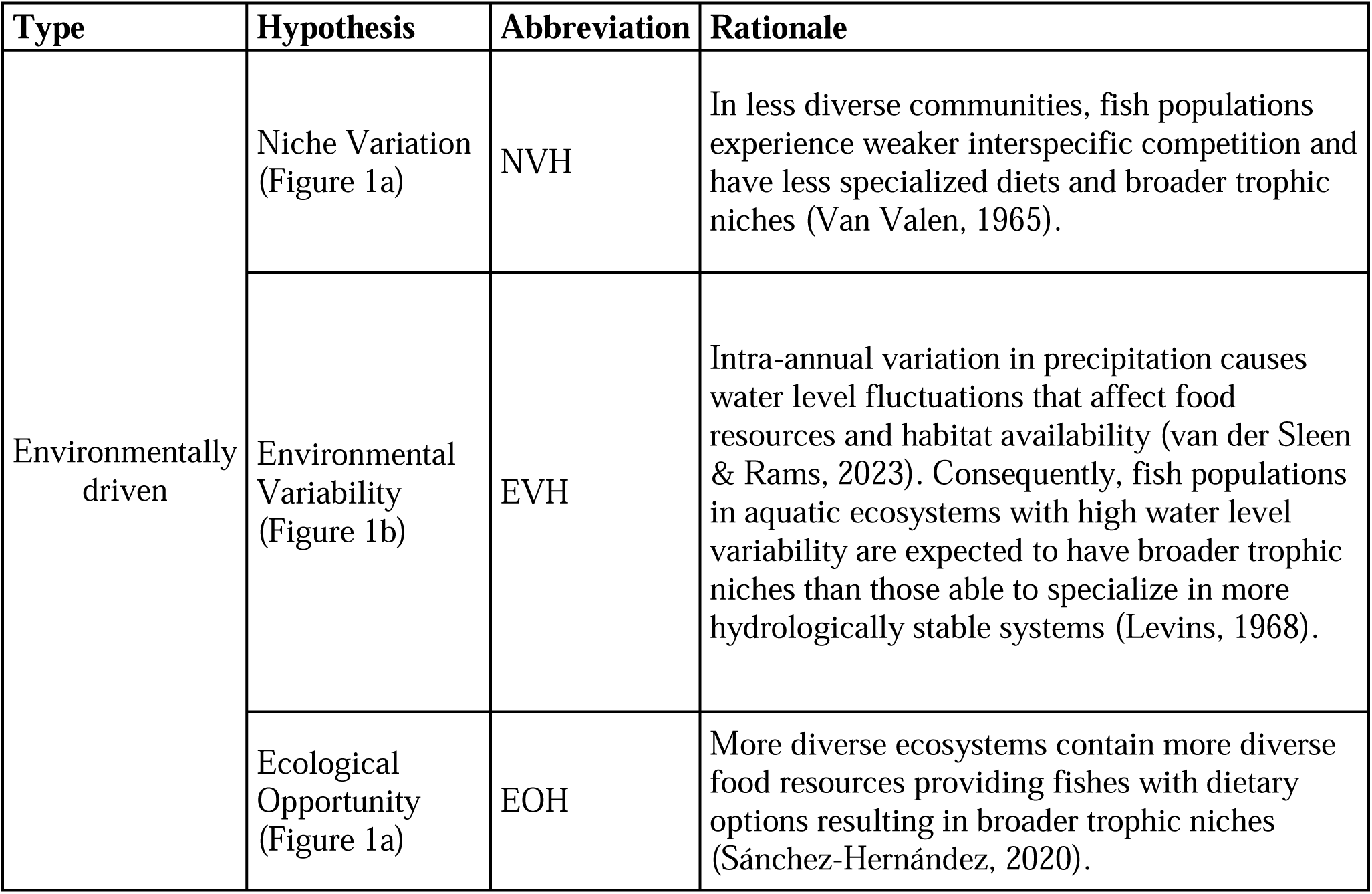
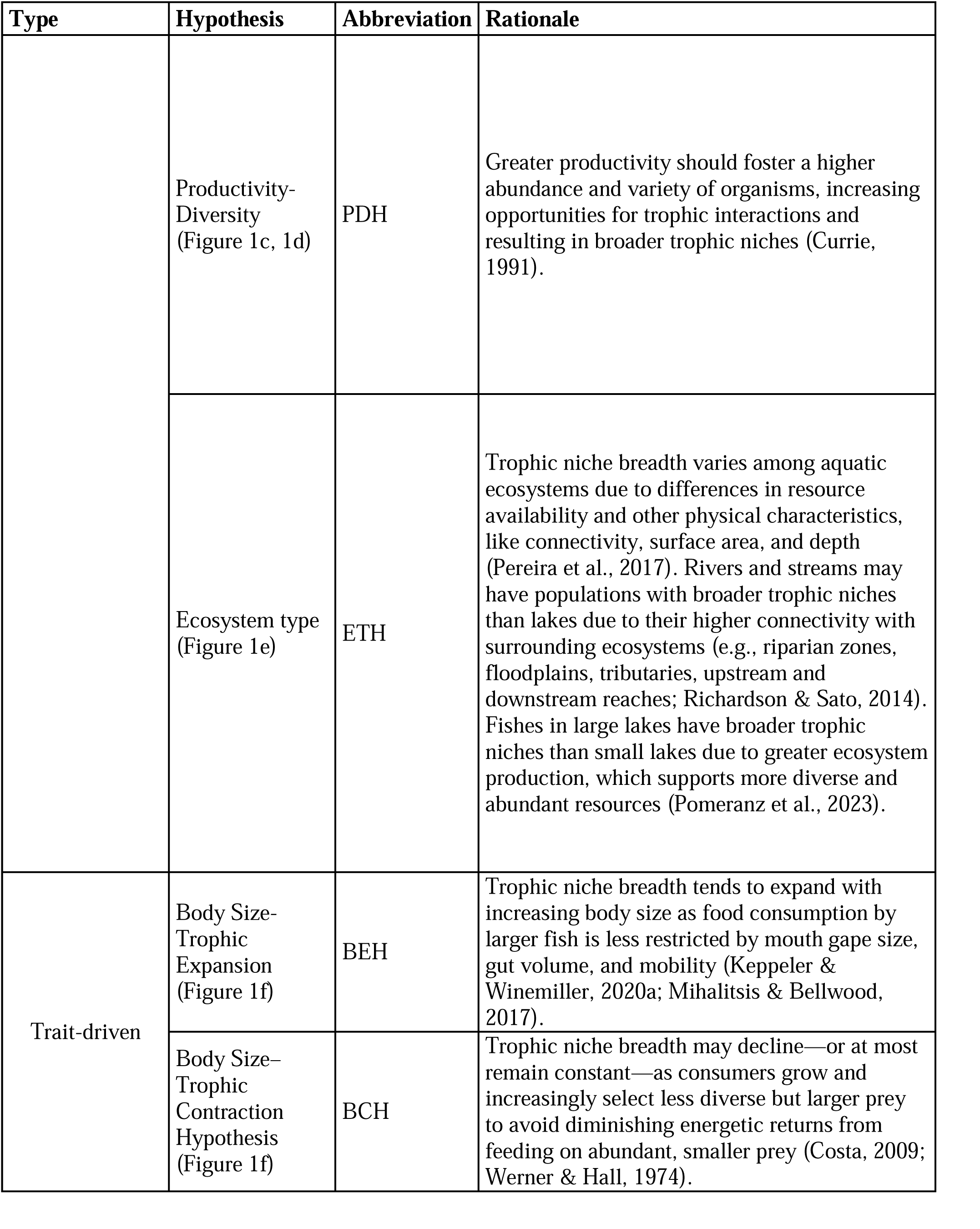
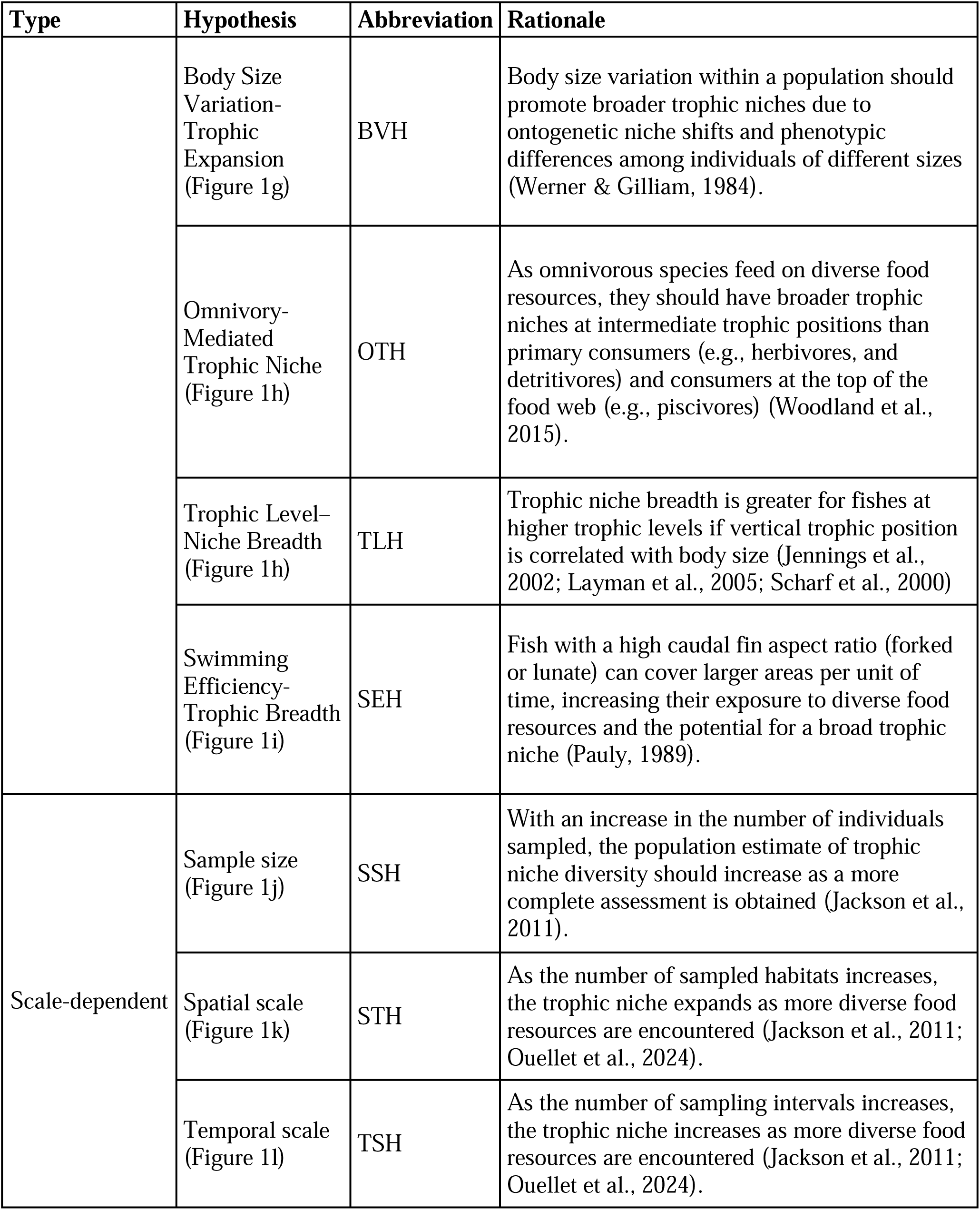
Description of existing and newly formulated trophic niche hypotheses tested with fish isospaces. Hypotheses and rationales are provided.

**FIGURE 1.**
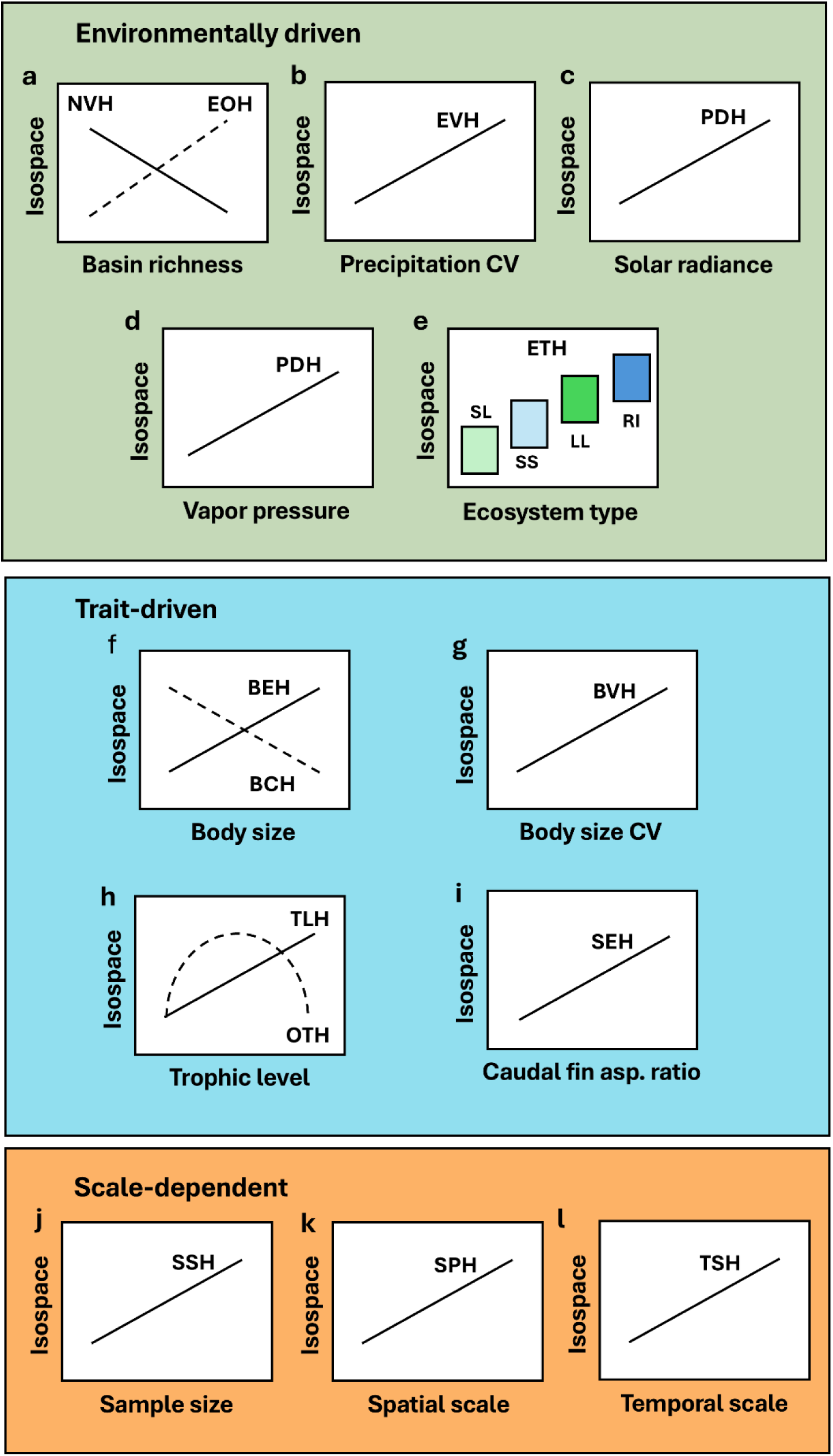
Theoretical associations between population isospace (δ¹ N and δ¹³C) and predictors related to environmental conditions (basin richness, coefficient of variance [CV] of precipitation, solar irradiance, vapor pressure, and ecosystem type), species traits (average body size, CV of body size, trophic level, and caudal fin aspect ratio), and scale (sample size, spatial scale, and temporal scale). For graphical simplicity, trophic level is presented instead of the trophic guild. Low trophic positions correspond to primary consumers (herbivores-frugivores and algivorous-detritivores), intermediate trophic positions to omnivores, invertivores, and planktivores, and high trophic positions to carnivores and piscivores. Lines, curves, and boxes portray hypothesized relationships (see Table 1 for the rationales). Hypotheses: NVH – Niche Variation, EVH – Environmental Variability, EOH – Ecological Opportunity, PDH – Productivity-Diversity, ETH – Ecosystem Type, BEH – Body Size-Trophic Expansion, BCH – Body Size–Trophic Contraction, BVH – Body Size Variation-Trophic Expansion, OTH – Omnivory-Mediated Trophic Niche, TLH – Trophic Level–Niche Breadth, SEH – Swimming Efficiency-Trophic Breadth, SSH – Sample Size, STH – Spatial Scale, TSH – Temporal Scale. Ecosystem type: SL – Small lake, SS – Small stream, LL – Large lake, RI – River.

Among the environmentally driven hypotheses, those related to latitudinal gradients are particularly relevant due to their implications for biodiversity patterns and climate change responses (De Frenne et al., 2013). Latitudinal gradients in the degree of temporal variation in environmental conditions have long been proposed to influence niche breadth (Dobzhansky, 1950). Lower seasonal environmental variability in the tropics is expected to lead to more stable populations with narrower niches (Vázquez & Stevens, 2004). Interspecific competition is also expected to be stronger in tropical communities that harbor higher species diversity, which promotes niche partitioning, specialization, and diversification that in turn facilitates species packing (Niche Variation Hypothesis – NVH; Pellissier et al., 2018; Van Valen, 1965) (Figure 1a). To date, evidence for narrower trophic niches at lower latitudes is limited, weak, and inconclusive (Cirtwill et al., 2015), and the prediction may not apply to some ecosystems. For example, dynamic hydrology may promote broad trophic niches in both temperate and tropical rivers (Environmental Variability Hypothesis – EVH; Levins, 1968; van der Sleen & Rams, 2023) (Figure 1b).

The larger geographic area and the greater solar radiation in the tropics enhance productivity, expanding the available trophic niche space. Greater and more stable ecosystem productivity in the tropics might support greater biodiversity without necessarily constraining population trophic niches (Brown, 2004; Davies et al., 2007). The Ecological Opportunity Hypothesis (EOH; Figure 1a) suggests that when food resources are abundant, greater species diversity can lead to broader trophic niches (Pereira et al., 2017; Sánchez Hernández et al., 2020). However, not all tropical and subtropical regions are highly productive. Arid regions— such as the African Sahel, Brazilian Caatinga, Northern Australia, and the Arabian Peninsula— have hot, dry conditions that limit ecosystem productivity and diversity (Hawkins et al., 2003), which, in turn, may limit trophic niche breadth (Productivity-Diversity Hypothesis - PDH; Currie, 1991) (Figure 1c, 1d).

The characteristics of the ecosystems in which populations are embedded likely play a crucial role in defining their trophic niche (Ecosystem Type Hypothesis – ETH) (Figure 1e). For example, closed or semi-closed systems, such as lakes, limit species’ ability to disperse and encounter diverse prey (Pereira et al., 2017). Larger ecosystems, like high-order rivers and great lakes, generally support more extensive and complex food webs due to higher total productivity, diversity and stability (Pomeranz et al., 2023). Conversely, smaller ecosystems, such as headwater streams, wetlands, and ponds, receive large resource subsidies from adjacent terrestrial ecosystems (Richardson & Sato, 2014; Vannote et al., 1980), fostering populations with broad trophic niches.

Species traits are increasingly recognized as key determinants of species’ trophic niches, and trait-driven hypotheses have become a common framework for explaining niche variation (Keppeler & Winemiller 2020b; McGill et al., 2006). Body size has gained a particular focus owing to its ease of measurement, large natural variation, and strong associations with metabolism, energy demand, longevity, foraging area, food web structure, and other physiological and ecological factors (Arim et al., 2010; Rooney et al., 2008; Woodward et al., 2005). The Body Size-Trophic Expansion Hypothesis (BEH; Figure 1f) suggests that as fish grow larger, they can consume a wider range of prey sizes, particularly if they are gape-limited predators (Keppeler & Winemiller, 2020b; Mihalitsis & Bellwood, 2017). Conversely, the Body Size–Trophic Contraction Hypothesis (BCH; Figure 1f) posits that organisms optimize their diets by becoming more selective as they grow, focusing on larger, more profitable prey while avoiding abundant, smaller prey (Costa, 2009; Werner & Hall, 1974). Populations composed of individuals with different body sizes are likely to have broader trophic niches (Body Size Variation-Trophic Expansion Hypothesis - BVH) (Figure 1g) due to ontogenetic niche shifts (Nakazawa, 2014; Werner & Gilliam, 1984) that increase between-individual variation in size-structured populations (Bolnick et al., 2003).

The relationship between trophic niche breadth and body size may differ among trophic guilds. For example, the vertical trophic position of predatory fishes tends to increase with body size and the average size of consumed prey, whereas detritivorous fishes show little variation in trophic position, and trophic niche breadth relative to body size (Keppeler et al., 2020; Keppeler & Winemiller, 2020b). Trophic niche breadth may increase with trophic position (Trophic Level– Niche Breadth -TLH) as predatory fishes, with their large mouths, are capable of ingesting a relatively broad spectrum of prey compared to fishes at lower trophic levels (Jennings et al., 2002; Layman et al., 2005; Scharf et al., 2000) (Figure 1h). Conversely, the Omnivory-Mediated Trophic Niche Hypothesis (OTH; Figure 1h) suggests that trophic niche breadth varies predictably with vertical trophic position, being greatest for omnivorous species at intermediate levels due to higher proclivity for foraging across multiple trophic levels (Woodland et al., 2015).

Mobility could influence trophic niche breadth if more vagile organisms can access and exploit a greater variety of habitats and food resources (Pauly, 1989). In fish, the shape of the caudal fin is strongly associated with patterns of locomotion and vagility. Species with forked or lunate caudal fins are typically continuous, fast swimmers, whereas those with broad, truncated, or rounded caudal fins tend to be slower swimmers with less endurance but greater lateral maneuverability (Sambilay, 1990). Fish having a caudal fin with a high aspect ratio (forked, lunate) may cover a larger area per unit of time (Pauly, 1989) and thereby exploit more diverse food resources (Rooney et al., 2008), resulting in a positive relationship between fin aspect ratio and trophic niche breadth (Swimming Efficiency-Trophic Breadth Hypothesis – SEH) (Figure 1i).

Niche breadth estimates based on large numbers of individuals are likely higher than those based on small samples (Sample Size Hypothesis – SSH; Jackson et al., 2011) (Figure 1j). Estimates of niche breadth are likely to be further inflated if the sample includes individuals from different locations and habitats (Spatial Scale Hypothesis – STH; Figure 1k) or periods (Temporal Scale Hypothesis – TSH; Figure 1l) (Ouellet et al., 2024). Failure to account for these factors may bias estimates used in comparative studies of trophic ecology, particularly regarding niche breadth (Jackson et al., 2011).

Given the multiple factors that may influence trophic niche breadth estimates, comparative studies at large spatiotemporal scales are rare (e.g., Franco-Trecu et al., 2022; González et al., 2023; Keppeler et al., 2020a). Most comparative studies of the trophic niche have examined population mean differences without explicit analysis of intraspecific variation (Bolnick et al., 2003; Des Roches, 2017). Another challenge is the lack of standardized methods for quantifying diet breadth among species that may have divergent foraging behaviors and associated morphologies. The usual methods used to characterize the diet of animals, such as stomach content analysis and fecal analysis, provide a snapshot of the diet and may not reflect patterns of food consumption over longer periods to provide a more reliable diet estimate. Furthermore, these conventional methods are generally limited in their ability to identify fragmented or partially digested material. They may not necessarily reflect foods assimilated by the individual, since hard food items tend to have a longer retention time in the gut and are digested less efficiently than soft items (Araújo et al., 2007; Keppeler et al., 2020b; Votier et al., 2003).

Stable isotope analysis (SIA) has emerged as a useful tool to examine trophic niches variations in the 1970s (Bearhop et al., 2004; Layman et al., 2012, Turner et al., 2023). Since then, ratios of ^15^N/^14^N (δ^15^N) and ^13^C/^12^C (δ^13^C) of tissue samples have been increasingly used in food web research (Layman et al., 2012). δ^15^N has a positive correlation with trophic level due to gradual ^15^N enrichment (usually from 2–4‰, termed “trophic fractionation”) during the assimilation of ingested material into consumer tissue (Post, 2002). Variation in δ^13^C in plants and other primary producers is associated with, among other factors, differences in the photosynthetic pathways (i.e., C3, C4, CAM). These differences in isotopic ratios of C and other elements provide a basis for inferring animal assimilation of biomass derived from primary producers using mixing models that integrate isotopic signatures of multiple potential sources (Fry, 2007; Parnell et al., 2013; Stock et al., 2018). Some advantages of SIA are the ability to obtain data from small tissue amounts (in some cases tissue samples can be obtained by non-lethal methods; e.g., Fredrickson et al., 2021; Maitland & Rahel, 2021) and the ability to estimate dietary histories over varying time intervals by analyzing incremental tissues (e.g., otoliths), tissues with different turnover rates or samples collected during various periods (Newsome et al., 2009).

We compiled an extensive database of stable isotope signatures (δ^13^C, δ^15^N) of freshwater fishes to perform a global analysis of relationships between population isotopic space (referred to as “isospace”) and variables associated with climate, biodiversity, ecosystem type, species traits, and sampling effort. We used solar radiation as a metric of energy availability (Brown et al., 2004), precipitation variation as a proxy of water level fluctuations and seasonality (Oki et al., 2004), and vapor pressure as an indicator of temperature and humidity, which is particularly useful for distinguishing tropical rainforests from drier and/or colder ecosystems (Kottek et al., 2006). In addition, we included 1) basin richness (i.e., number of fish species) as a proxy for potential, either current or past, interspecific competition (Connell, 1980); 2) aquatic ecosystem type as a basic descriptor of aquatic habitat conditions (e.g., spatial extent, connectivity) influencing food web structure (Riede et al., 2010); 3) average body size as a trait affecting food consumption (Keppeler & Winemiller, 2020a); 4) variation in body size as a metric reflecting the potential for ontogenetic trophic niche shifts (Werner & Gilliam, 1984); 5) trophic guild as a description of feeding preferences (Parravicini et al., 2020); and 6) caudal fin aspect ratio as a proxy for potential mobility and spatial extent of foraging activity (Pauly, 1989). To account for differences in study design and survey methods, we analyzed three metrics related to sampling scale: the number of samples, the number of years sampled, and the diversity of habitats sampled. Our overarching goal was to test if population isospaces provide support for the aforementioned hypotheses of trophic niche variation (Table 1, Figure 1).

## METHODS

### Stable Isotope Values and Species Traits

We compiled a dataset including 33,509 values of δ^13^C and δ^15^N obtained from muscle tissue of 1,422 populations of 880 freshwater fish species from six major ecoregions across the globe (Figure 2; Appendix S1, Table S1). We use ‘population’ as a pragmatic term in this study, referring to fish species sampled from a distinct water body. δ¹ N represents the ratio of ¹ N to ¹ N in a sample relative to atmospheric nitrogen, and δ¹³C represents the ratio of ¹³C to ¹²C relative to the Pee Dee Belemnite standard. Both values were expressed in parts per thousand (‰). We did not apply mathematical lipid corrections to δ¹³C values prior to analysis, as recommended by Post et al. (2007) for two main reasons. First, C:N ratios were unavailable for some datasets; second, recent studies have shown that lipid corrections are tissue- and species-specific, with no universally reliable C:N ratio threshold (Cloyed, 2019).

**FIGURE 2.**
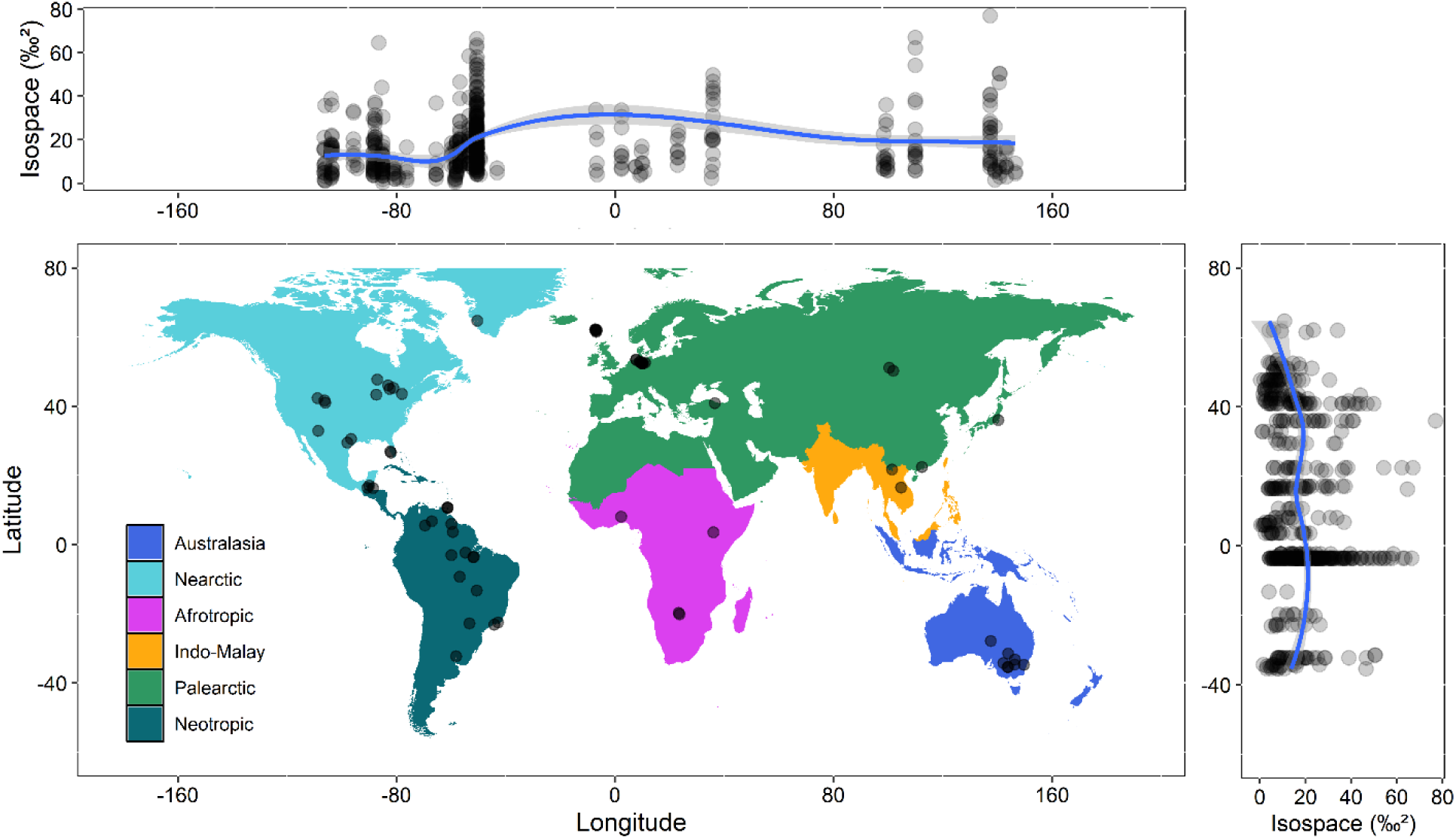
Spatial variation in isospace size. The upper panel illustrates variation across longitude, and the right panel depicts the relationship across latitude. Trend lines are fitted using locally estimated scatterplot smoothing (LOESS). Points on the map represent the locations of the studied water bodies.

The isotopic data were derived from studies conducted by the authors and based on sample collections representative of the local fish community, including multiple species from different trophic guilds and species with robust sample sizes (Appendix S1, Table S1). The data were then filtered to include only populations with complete body size information for all individuals and with at least 10 individuals sampled. The studies also provided information about sampling dates, locations, and habitats.

Individual body sizes were measured either with body length (standard length or total length, mm) or body mass (g) (Appendix S2, Table S1). To standardize, we converted all body lengths to body mass using the allometric formula (Keys, 1928):

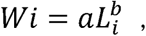

where *Wi* is the predicted weight of individual *i*, *Li* is its length, and log(*a*) and *b* represent the intercept and slope, respectively, in the logarithmic form of the length-weight relationship for the population of individual *i*. The parameters *a* and *b* for each species were estimated using posterior modes derived from kernel density estimation, following the Bayesian hierarchical approach proposed by Froese et al. (2014), and obtained via the R package *fishflux* (Schiettekatte et al., 2020). Then, the average body size along with their respective coefficients of variation were calculated for each fish population.

The caudal aspect ratio (AR) of each species was obtained from the *FishLife* R package (Thorson, 2023) and is calculated as follows (Pauly, 1989):

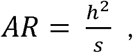

where *h* is the height of the caudal fin and *s* is the surface area of the caudal fin. Low aspect ratios (<1.5) are associated with truncate and rounded fins; intermediate aspect ratios (1.5–3) are associated with forked and emarginate fins, and high aspect ratios are associated with lunate fins (>3) (Table S3) (Sambilay, 1990).

Fishes were classified into eight trophic guilds based on diet studies (Appendix S2, Table S1): 1) Algivore/Detritivore, 2) Herbivore/Frugivore, 3) Omnivore (consume primary producers and invertebrates), 4) Planktivore, 5) Invertivore, 6) Carnivore (commonly consume macroinvertebrates, fish, and/or other vertebrates), 7) Piscivore, and 8) Fish Parasite (blood, mucus, and scale feeders). When diet studies were unavailable for a given species, we inferred its trophic guild based on closely related species (i.e., within the same genus, or rarely, family). Given that juvenile and adult size classes often show dietary differences, our classification is based on adult individuals.

### Environmental variables

Monthly averages of solar radiation (kJ m ² day ¹), precipitation (mm), and vapor pressure (kPa) from 1970 to 2000 were obtained at a spatial resolution of 0.5 arcminutes for the entire globe using the WorldClim Version 2 database (Fick & Hijmans, 2017) via the *geodata* R package (Hijmans et al., 2024). The WorldClim database captures substantial climatic variability (annual averages ranging from −23 °C to 29 °C, depending on the region; Mourshed, 2016). Recent climate change (∼1 °C increase in global mean temperature over the past two decades; NASA, 2023) introduces a potential source of error; however, this is not expected to bias our analyses and conclusions as the degree of change is minimal in the entire range of temperatures in the data set.

We used the compiled climate databases to calculate average annual solar radiation, annual coefficient of variation of precipitation, and average annual vapor pressure for each data location (Appendix S3, Figure S1). For basin richness, we estimated the number of fish species in each sub-basin where the stable isotope studies were conducted. These biodiversity estimates (Appendix S3, Table S1) were based on values obtained from the literature or regional surveys reported by the studies producing the isotopic data included in our analyses.

We classified the sampled water bodies into four main categories: large lakes (>150 km²), small lakes (<150 km²), streams (river order 1–3), and rivers (river order >3). As the populations within each ecosystem type could inhabit more than one habitat, for each population we calculated the Shannon diversity of habitat types from which fish individuals were collected (i.e., 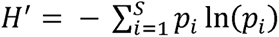, where S is the total number of habitats, *p_i_* is the proportion of individuals in the *i*th habitat, and ln(*p_i_*) is the natural logarithm of the proportion) (herein referred as habitat diversity). To do this, we classified sampling habitats into four broad categories to align with the dataset’s resolution: main river channel, stream, lake, and other floodplain habitats (e.g., flooded forest, temporary pools). We decided to use the habitat diversity rather than the number of habitats sampled due to the higher predictive performance of the former. Similarly, we recorded the number of years that each population was sampled. In this case, the number of years sampled had better predictive performance than the diversity of years from which fish individuals were collected.

We classified the geographic locations from where tissues were collected into six major ecoregions: 1) Neotropic, 2) Palearctic, 3) Indo-Malay, 4) Afrotropic, 5) Nearctic, and 6) Australasia (Olson et al., 2001). We also classified the sampling according into four main climate zones: Tropical (0°–23.5° N/S), Sub-tropical (23.5°–35° N/S), Temperate (35°–60° N/S), and Polar (60°–90° N/S) (Peel et al., 2007). Although ecoregions and climate zones do not directly affect our hypotheses, their inclusion provided additional spatial variables for comparison of fish isospaces and description of data structure.

### Data analysis

We estimated the isospace of each fish population using the kernel utilization density method proposed by Eckrich et al. (2019) (Appendix S4). We chose the kernel utilization density because, unlike the standard ellipse area approach (Jackson et al., 2011), it does not assume a normal distribution in two-dimensional space and is less sensitive to outliers than the convex hull method (Layman et al., 2007). We defined the kernel utilization density isospace at 50%, 75%, and 90% contours for each population. Since these contours were highly autocorrelated (Pearson correlation >0.97), only the 75% contour was used for isospace modeling (see below).

To ensure a more robust and representative assessment, we estimated isospace only for fish populations with at least 10 individuals. Overall, 541 fish populations from 358 species met the minimum requirements of a sample size of at least 10 with accompanying body size measurements. Sensitivity tests were conducted to evaluate whether increasing this threshold to 15 individuals, as suggested by Eckrich et al. (2019), would alter the results. While the overall patterns remained consistent, increasing the threshold led to the removal of 108 populations, weakening the observed trends. Species classified as fish parasites were removed from analyses due to the small number of populations in the database (N = 4).

Three approaches were used to investigate the drivers of fish population isospace variation: multiple regression (MR), phylogenetic generalized least squares (PGLS), and boosted regression trees (BRT). MR is a comparatively simpler model assuming that population responses to predictors are independent and linear and serving as a baseline for comparison with PGLS and BRT results. PGLS accounts for non-independence of populations due to shared ancestry, providing estimates of relationships between variables within an evolutionary context (Freckleton et al., 2002). Since a complete phylogenetic tree for fishes is not yet available, shared ancestry was estimated using taxonomic information retrieved from the Integrated Taxonomic Information System (ITIS; NMNH, 2025) via the *taxize* R package (Chamberlain et al., 2020). The taxonomic signal of population isospace was quantified using Pagel’s lambda (Pagel, 1999) through the *phytools* R package (Revell, 2024). BRT is an ensemble machine-learning algorithm that combines multiple regression trees to enhance predictive accuracy. BRT employs an iterative learning process in which each tree is fitted to the residuals of the previous tree, sequentially reducing errors (Elith et al., 2008). While BRT does not account for non-independence of populations due to shared ancestry, it is a powerful and flexible algorithm capable of handling non-linear relationships, interactions, and multicollinearity (Elith et al., 2008; Pichler & Hartig, 2023). BRT models were constructed using learning rates of 0.001, tree complexity of 5, bag fraction of 0.6, and Gaussian family distribution (Elith et al., 2008).

We tested five model structures for all model approaches (i.e., MR, PGLS, and BRT). The scale-dependent model included sample size, habitat diversity, and the number of sampling years as predictors. The trait-driven model incorporated average body size, body size CV, trophic guild, and caudal fin aspect ratio. The environmentally driven model used average solar radiance, precipitation CV, average vapor pressure, ecosystem type, and basin fish richness as predictors. The full model combined all predictors from the previous three models, while the intercept-only model (i.e., *Y* = β_0_ +*∊*, where β_0_ is simply the mean value of the response variable across all observations) contained no predictors. To compare model performance, we used three main metrics: i) the Akaike Information Criterion (AIC) that balance model complexity with goodness-of-fit; 2) explanatory power (*R²-explanatory*) to assess in-sample error; and 3) predictive power (*R²-predictive*) based on five-fold cross-validation to evaluate out-of-sample error.

We performed backward model selection to simplify the best BRT model following the approach of Elit et al. (2008) that relies on 5-fold cross-validation and the change in predictive deviance to identify the best model. We compared the performance of the best BRT model with the best PGLS and MR models using exploratory and predictive power (5-fold cross-validation). To interpret the relationship between predictors and fish isospace estimated for BRT, we used partial dependence plots revealing this relationship after accounting for the average effects of all the other model predictors (Friedman, 2001; Friedman & Meulman, 2003). Locally Estimated Scatterplot Smoothing (LOESS) was performed for the plots to better visualize the main trends. In addition, LOESS was used to describe latitudinal and longitudinal gradients of fish isospaces.

Given concerns about isospace compression as resource pathways become integrated at higher trophic positions (Hoeinghaus & Zeug, 2008), we also conducted BRT models separately for predators (Invertivores, Carnivores, and Piscivores) and non-predators (Algivores/Detritivores, Herbivores/Frugivores, Omnivores, and Planktivores). We opted to analyze these two broad groups rather than individual trophic guilds due to the small sample sizes and limited spatial distributions of some guilds (e.g., herbivores being largely confined to tropical regions), which could bias interpretations of environmental gradients. In addition, previous studies have shown that predators and non-predators may respond differently to environmental gradients and may differ in how traits influence trophic niche and trophic position (Keppeler et al., 2020).

All analyses were carried out in R, version 4.4.1 (R Core Team, 2024). BRT was carried out in the *dismo* package (Hijmans et al., 2023), PGLS in the *caper* package (Orme et al., 2023), and MR and LOESS in the *stats* package (R Core Team, 2024). The *MuMIn* package was used for model selection with AIC (Bartoń, 2024), and the *caret* package was used for cross-validation (Kuhn, 2008).

## RESULTS

### Dataset composition

The tropics were the most represented region in the dataset, with 328 fish populations, followed by temperate (154), subtropical (62), and polar regions (7). All fish populations from the polar zone were carnivores. In the temperate zone, approximately 64% of fish populations were predators (invertivores, carnivores, or piscivores). This proportion declined to 52% in the subtropics and 45% in the tropics. Omnivores were more prevalent in the subtropics (35%) and tropics (34%) than in the temperate zone (27%). Similarly, algivores/detritivores and herbivores/frugivores were more common in the tropics (19%) and subtropics (11%) compared to the temperate region (4%). Planktivores were more frequent in the temperate zone (∼5%) but were less represented in other regions (<2%).

The Neotropics were the most represented ecoregion, with 313 fish populations, followed by the Nearctic (105), Palearctic (66), Australasia (25), and both the Afrotropic and Indo-Malay ecoregions (21 each). The highest proportion of predators was observed in the Afrotropic biome (86%), followed by the Palearctic (66%), Australasia (64%), and Nearctic (63%). Omnivores were most prevalent in the Indo-Malay (43%) and Neotropics (36%). Algivores/detritivores and herbivores/frugivores were more prominent in the Neotropics (20%), Australasia (12%; exclusively algivores/detritivores), and Indo-Malay (10%). Planktivores were more frequent in the Palearctic (8%), Afrotropic (5%), and Nearctic (4%). Additional details are available in Appendix S5.

### Isospace variation across space and fish families

We observed large variation in population isospace size across geographic regions (Figure 2, 3a) and fish families (Figure 3b). Overall, population isospaces were smaller in the Nearctic and Australasian ecoregions, and largest in the Neotropics, Palearctic, and Indo-Malay (Figure 3a). Furthermore, a consistent latitudinal gradient in isospace size was not observed, although populations at latitudes higher than 40° tended to have smaller isospaces (Figure 2).

**FIGURE 3.**
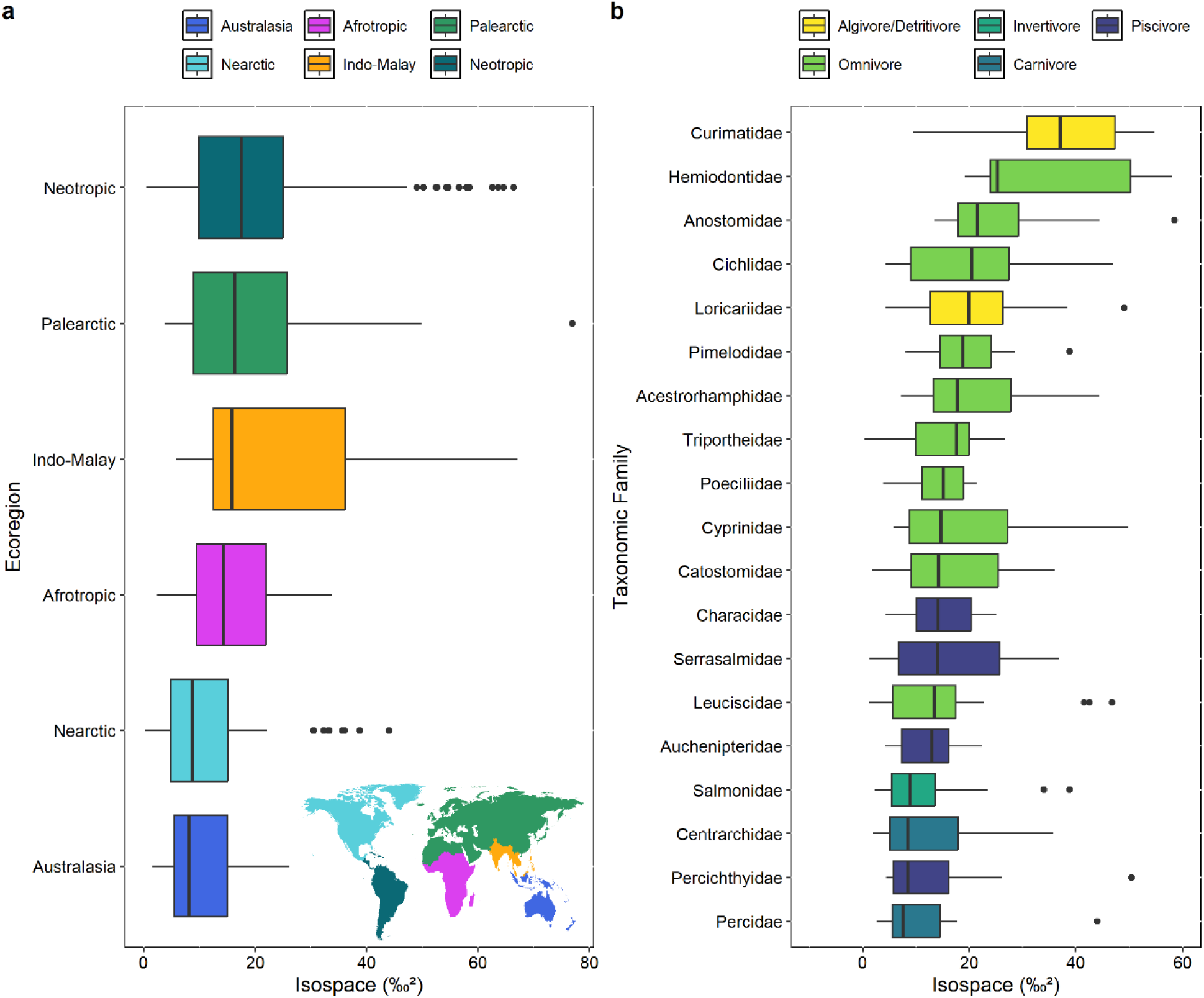
Isospace size of fish populations across ecoregions (a) and taxonomic families (b). Only those families with more than 10 populations analyzed are shown. Boxplot showing data distribution, including median, interquartile range (IQR), and outliers beyond the whiskers.

Neotropical algivorous-detritivorous fish in the families Curimatidae and Hemiodontidae exhibited the largest isospaces (Figure 3b). Conversely, families dominated by invertivorous and piscivorous species (e.g., Centrarchidae, Percidae, Percichthyidae, Salmonidae) from different ecoregions had considerably smaller isospaces. Isospace size had a significant taxonomic signal (Pagel’s lambda = 0.7, p < 0.001), indicating that closely related populations tend to have similar isospace sizes.

### Model comparisons and variable importance

Full models incorporating environmental, trait-dependent, and scale-dependent variables, performed best across MR, PGLS, and BRT models (Table 2). BRT models had superior explanatory (R^2^ = 0.77) and predictive power (R^2^ = 0.33) compared to MR and PGLS, indicating that the relationships between predictors and fish population isospace size are nonlinear with relatively minor interactions among predictors. The backward selection applied to the best BRT model (Full) did not eliminate any predictors, suggesting that all variables contribute meaningfully to predicting population isospace size. Accordingly, we describe the relationships between predictors and population isospace size as estimated by the Full BRT model.

**TABLE 2.** Comparison of results from three modeling approaches: multiple regression (MR), phylogenetic generalized least squares (PGLS), and boosted regressions tree (BRT) used to explain and predict variation in fish population isospace. Models were ranked according to their predictive capacity (R^2^ Pred). AIC was not calculated for BRT models due to the inherent limitation on quantifying model complexity.

| Model structure | $R^2$ Pred<br>(Cross-validation) | $R^2$ Expl | AIC | Delta | Weight |
| --- | --- | --- | --- | --- | --- |
| **MR** |  |  |  |  |  |
| Full | 0.20 (0.10) | 0.25 | 4200 | 0 | 1.00 |
| Scale | 0.14 (0.05) | 0.14 | 4242 | 43 | 0.00 |
| Traits | 0.11 (0.06) | 0.13 | 4262 | 61 | 0.00 |
| Environment | 0.08 (0.04) | 0.09 | 4281 | 81 | 0.00 |
| Intercept-only | 0.00 (0.00) | 0 | 4317 | 116 | 0.00 |
| **PGLS** |  |  |  |  |  |
| Full | 0.21 (0.05) | 0.21 | 4119 | 0.0 | 1.00 |
| Environment | 0.13 (0.05) | 0.12 | 4171 | 51.9 | 0.00 |
| Scale | 0.12 (0.02) | 0.13 | 4143 | 24.0 | 0.00 |
| Traits | 0.07 (0.04) | 0.06 | 4193 | 73.3 | 0.00 |

| Model structure | R<sup>2</sup> Pred<br>(Cross-validation) | R<sup>2</sup> Expl | AIC | Delta | Weight |
| --- | --- | --- | --- | --- | --- |
| Intercept-only | 0.00 (0.00) | 0.00 | 4210 | 90.7 | 0.00 |
| **BRT** |  |  |  |  |  |
| Full | 0.33 (0.00) | 0.77 | - | - | - |
| Environment | 0.20 (0.00) | 0.27 | - | - | - |
| Traits | 0.16 (0.00) | 0.37 | - | - | - |
| Scale | 0.16 (0.00) | 0.26 | - | - | - |
| Intercept-only | 0.00 (0.00) | 0.00 | - | - | - |

Together, traits accounted for a cumulative relative contribution of 44%, while scale and environmental variables each contributed approximately 28%. Among the analyzed traits, trophic guild was the strongest predictor of isospace size (Relative Contribution [RC] = 18%), with body size CV (RC = 11%) and caudal fin aspect ratio (RC = 10%) also emerging as important predictors. Among the predictors, vapor pressure ranked highest (RC = 10%), with solar radiation (RC = 6%) and basin richness (RC = 6%) also showing high importance. Of the scale-dependent variables, habitat diversity of survey locations was the strongest predictor (RC = 11%), followed by number of years sampled (RC = 10%) and sample size (RC = 7%) (Figure 4).

**FIGURE 4.**
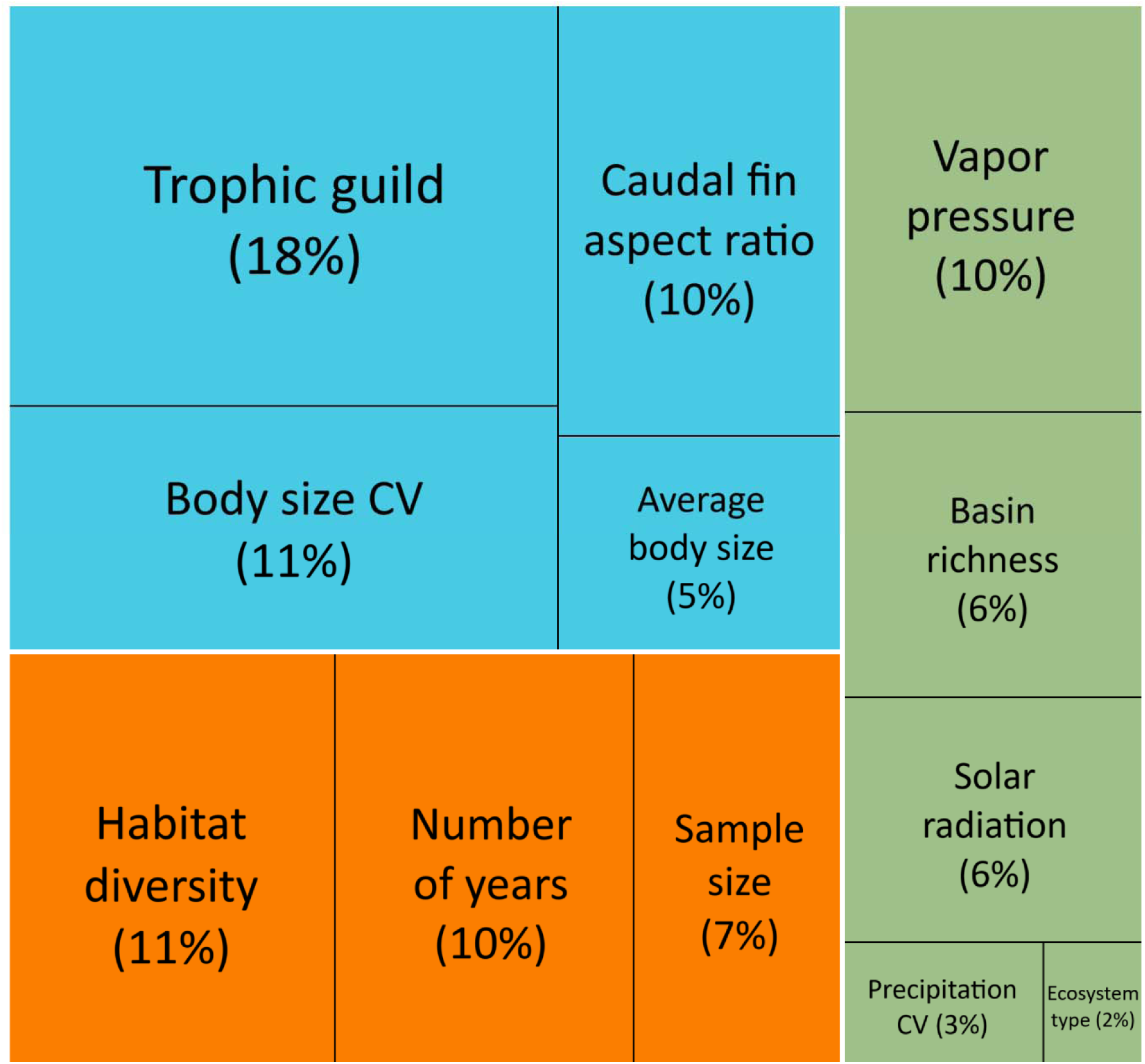
Relative contributions of predictor variables in the Boosted Regression Tree Model (BRT) for fish population isospace size. The blue color indicates trait variables, green indicates environmental variables, and orange indicates scale-dependent variables.

### Isospace variation — Environment

Population isospace size was negatively associated with basin richness, with a sharp decline of ∼13% from areas with less than 50 fish species (e.g., Greenland and Trinidad streams) to areas with 100 or more fish species (e.g., Uruguay River) (Figure 5a). Precipitation CV was positively associated with isospace size, albeit with a low magnitude of difference (∼5%) between regions with low variation in precipitation (e.g., rivers in southeastern Australia) and regions with high variation (e.g., rivers in the Okavango Delta, Botswana) (Figure 5b). The relationship between solar radiation and isospace size was weak and variable, with two peaks at intermediate and high radiation levels, where the isospaces were approximately ∼9–12% larger than those in low-radiation regions at higher latitudes (Figure 5c). Vapor pressure was positively associated with isospace size, with populations in cold regions, such as Lake Khövsgöl in Mongolia and streams in the northeastern U.S., having ∼33% smaller isospaces compared to those in tropical rainforest regions (Figure 5d). We observed relatively small differences (<4%) in isospace size across ecosystem types (Figure 5e). Fish populations in rivers had the largest isospaces, and those in large lakes (e.g., Lake Michigan) the smallest.

**FIGURE 5.**
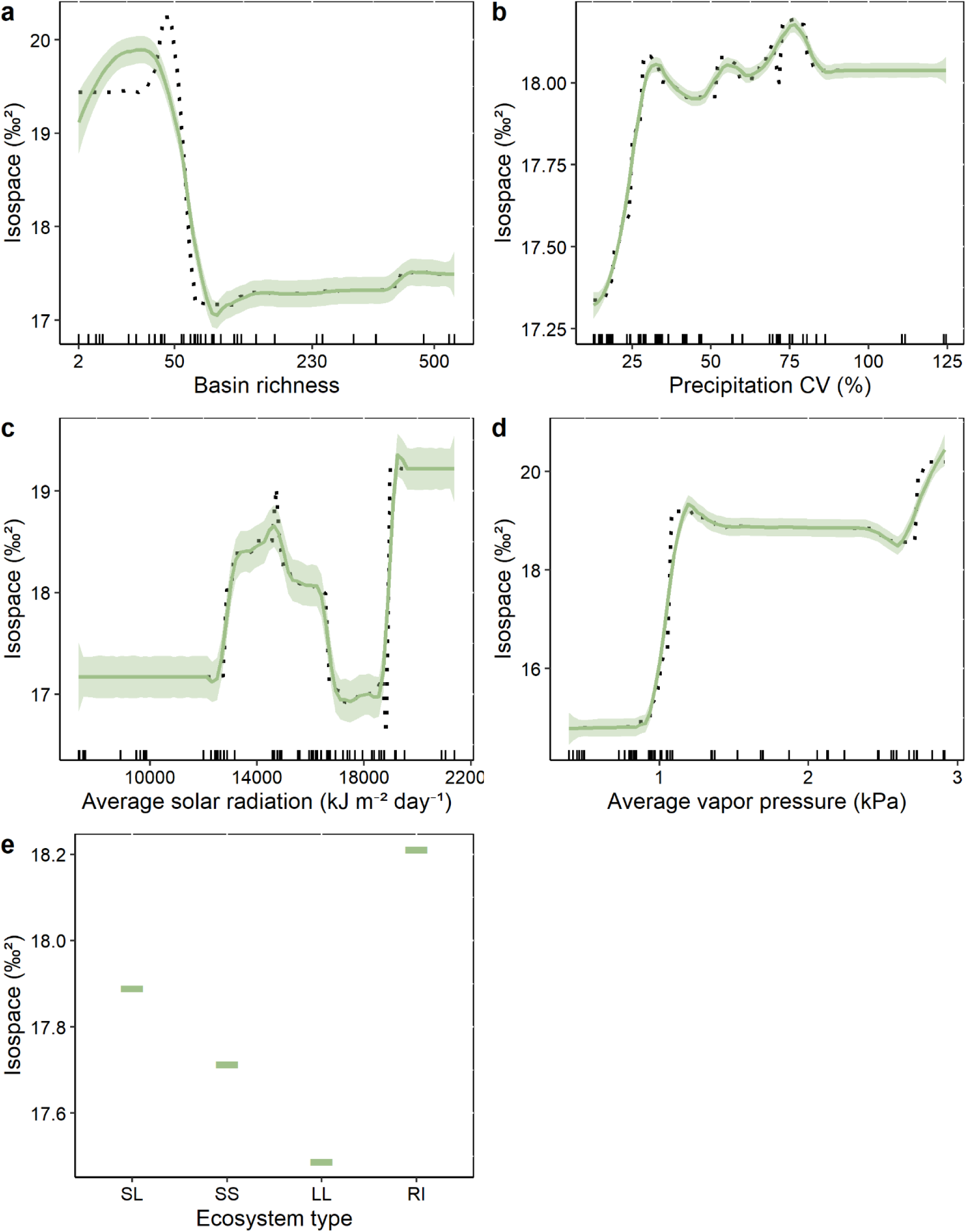
Marginal effects of five environmental variables on fish population isospaces. Dots and dashes are predictions based on the Boosted Regression Tree Model (BRT). Trend lines were fitted using locally estimated scatterplot smoothing (LOESS). SL= small lakes, SS = small streams, LL = large lakes, RI = rivers. The tick marks on the x-axes show the distribution of environmental values in the dataset.

### Isospace variation — Traits

The relationship between average body size and isospace size was weakly positive, with increases of 3% from fishes weighing 1 g to fishes weighing more than 5 kg (Figure 6a). High within-population variation in body size was also associated with larger isospaces, with an increase of approximately 35% along a gradient from 10 (i.e., the standard deviation is 10% of the mean) to 1000 in coefficient of variation (i.e., the standard deviation is ten times the mean value) (Figure 6b). Population isospace size differed significantly among trophic guilds, with Algivore/Detritivores and Herbivores/Frugivores having isospaces at least 26% larger compared to those of consumers at higher trophic levels (Figure 6c). Fishes with caudal fin aspect ratios <1.0 (truncate and rounded fins) had up to 22% larger isospaces than those with forked or emarginate fins (aspect ratio 1.5–3) and lunate fins (>3) (Figure 6d).

**FIGURE 6.**
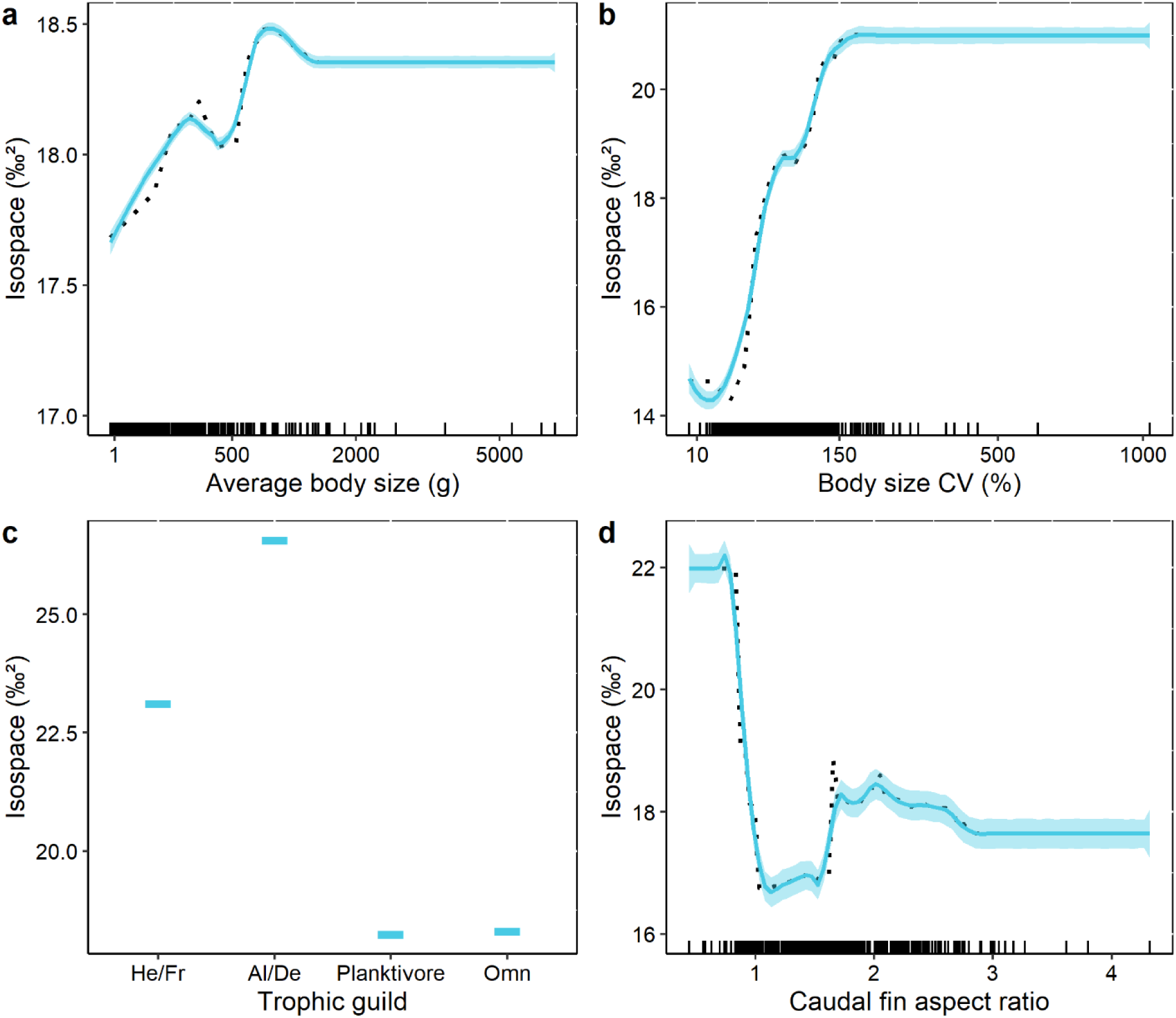
Marginal effects of four trait-related variables (a- average body size, b- the coefficient of variation [cv] of body size, c- trophic guild, and d- caudal fin aspect ratio) on the isospace of fish populations. Black dots are predictions based on the Boosted Regression Tree Model (BRT). Trend lines are fitted using locally estimated scatterplot smoothing (LOESS). The tick marks on the x-axes show the distribution of trait values in the dataset.

### Isospace variation — Scale

Population isospace size increased relative to spatial and temporal scales and sample size (Figure 7). Increasing the sample size from 10 to 100 was associated with an increase of up to 19% in fish population isospace size, whereas further increases from 100 to 600 resulted in little to no additional change (Figure 7a). An increase in habitat diversity from 0.1 (a single habitat) to 1 (three habitats with high equitability) was related to a ∼33% increase in isospace size (Figure 7b). Additionally, isospace size increased by ∼26% when the sampling period was extended from 1 to 9 years (Figure 7c).

**FIGURE 7.**
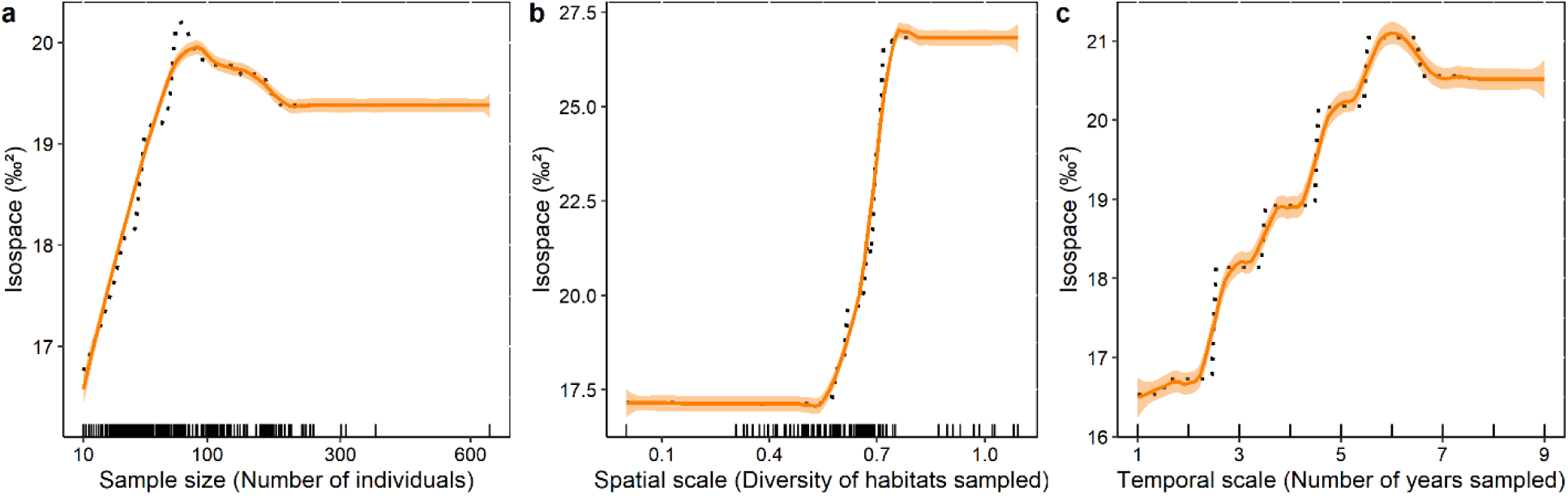
Marginal effects of sample size (a), and spatial (b) and temporal scales (c) on the isospace of fish populations. Black dots are predictions based on the Boosted Regression Tree Model (BRT). Trend lines are fitted using locally estimated scatterplot smoothing (LOESS). The tick marks on the x-axes show depict the distribution of habitat diversity (Shannon index) sampled, the number of years sampled, and the sample size values in the dataset.

### Isospace variation — Predators vs Non-Predators

BRT models performed separately for predators (invertivores, carnivores, and piscivores) and non-predators (algivores/detritivores, herbivores/frugivores, omnivores, and planktivores) revealed important differences. While predators followed trends similar to those observed when all trophic guilds were analyzed together, non-predators displayed distinct responses to four main variables. Among non-predators, isospace declined by ∼15% as average solar radiation rose from ∼12,000 to ∼19,500 kJ m ² day ¹ (Figure 8a), and expanded by ∼6% as basin richness increased from 7 to 540 species (Figure 8b). Isospace also increased by ∼4% as average body size increased from ∼1 g to ∼6.5 kg (Figure 8c), and by ∼5% in species with truncate or rounded caudal fins (caudal fin aspect ratio < 1.5) compared to those with lunate fins (caudal fin aspect ratio > 3) (Figure 8d). The relative contribution of predictors also shifted in non-predator models, with habitat diversity explaining the largest portion of variation (19%), followed by body size variation and trophic guild (14% each), sample size (13%), caudal aspect ratio (10%), average body size (8%), number of sampling years (6%), and average solar radiation (5%). All remaining predictors contributed less than 4%. Further details on predator and non-predator analyses are available in Appendix S6.

**FIGURE 8.**
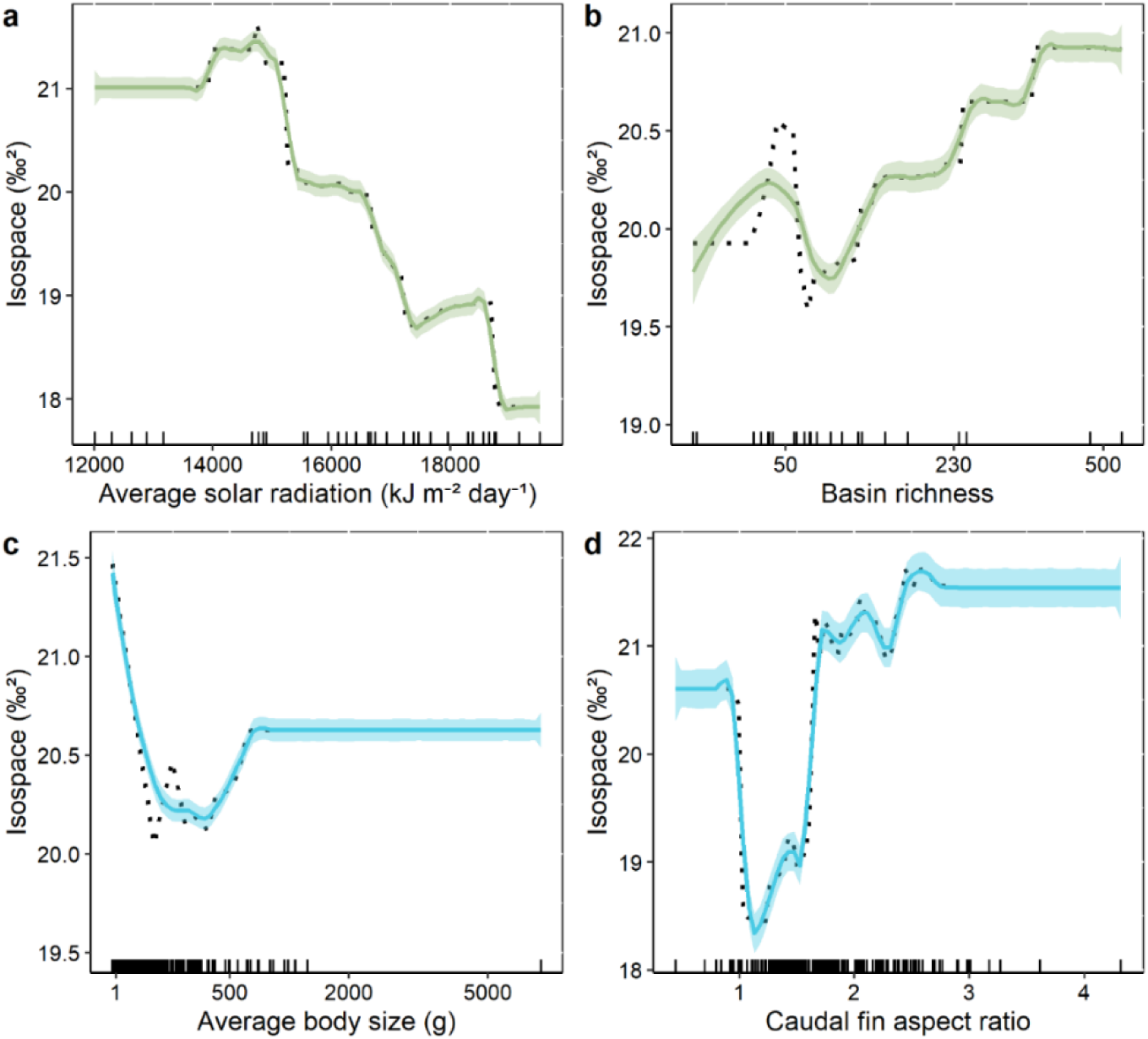
Marginal effects of two environmental variables (a – Average solar radiation, b- basin richness) and two traits (c- average body size, d - caudal fin aspect ratio) on the isospace of non-predatory fish populations. Black dots are predictions based on the Boosted Regression Tree Model (BRT). Trend lines are fitted using locally estimated scatterplot smoothing (LOESS). The tick marks on the x-axes show the distribution of trait values in the dataset.

## DISCUSSION

Our findings revealed considerable variation in fish population isospace size across the globe and taxonomic groups. The BRT model, which incorporated environmental, trait, and scale-dependent variables, explained up to 32% of the variation in isospace size during testing (i.e., out-of-sample predictions) — a relatively high predictive power for an ecological analysis performed at a global scale. The combined influence of environment, traits, and scale-dependent factors yielded the best predictive model, with environmental and trait-related variables being among the most important predictors. Our findings supported six of the eleven tested hypotheses linking trophic niche breadth with environmental and trait-associated mechanisms, providing moderate support for the Productivity-Diversity (PDH) and Body Size Variation-Trophic Expansion (BVH) hypotheses, and weak support for the Environmental Variability (EVH), Ecosystem Type (ETH), Niche Variation (NVH), and Body Size-Trophic Expansion (BEH) hypotheses. Of the five unsupported hypotheses (Ecological Opportunity [EOH], Omnivory-Mediated Trophic Niche [OTH], Body Size–Trophic Contraction [BCH], Trophic Level–Niche Breadth [TLH], and Swimming Efficiency-Trophic Breadth [SHE]), three were related to caudal fin aspect ratio and trophic guild, both of which showed consistent but opposite patterns to the predictions. Interestingly, the results differed between fish at the base and those at the top of the food web. While predators aligned with the patterns observed in the analysis encompassing all trophic guilds, non-predators showed considerable differences in four key variables—supporting the EOH, SHE and BCH hypotheses but diverging from the predictions of the PDH, and BEH. Isospace size variation exhibited scale dependence on both predators and non-predators, as suggested by the Sample Size (SSH), Spatial Scale (STH), and Temporal Scale (TSH) hypotheses.

### Isospace variation — Environment

Population isospaces tended to be larger in warm, humid regions, likely reflecting greater resource diversity and productivity, as proposed by the productivity–diversity hypothesis (PDH; Currie, 1991). In these environments, elevated energy availability and climatic stability may promote broader trophic opportunities and more consistent access to diverse resources. These conditions can support greater individual-level variation in resource use and trophic plasticity, ultimately contributing to population-level niche expansion. The greater availability of aquatic habitats in tropical and subtropical regions—partly driven by higher aquatic plant diversity and abundance—may further facilitate isotopic niche expansion. Aquatic plants provide structurally complex habitats that support diverse aquatic communities, including invertebrates that are important prey for many fish species (Thomaz & Da Cunha, 2010). In addition, co-occurrence of C3 and C4 plants is expected to be more frequent within hot, humid tropical regions, and the wide difference in δ^13^C between these plants at the base of aquatic foods could contribute to isotopic variation among consumers assimilating material derived from these sources (Sage & Monson, 1999; Still et al., 2003). This contrasts with cold regions, where low temperatures reduce metabolic rates, ecosystem productivity, and food resource diversity (Brown et al., 2004). Similarly, dry environments often cause water bodies to shrink or dry completely for intervals that restrict fish movement and foraging opportunities (Williams, 2006), degrade water quality (Missaghi et al., 2017), and limit basal resource production (Jaleel et al., 2009).

Supporting the NVH hypothesis, fish isospace size was negatively related to basin richness, but this relationship only emerged after accounting for climate variables associated with humidity and temperature (i.e., after including both in the model), which were positively correlated with isospace size. This suggests that resource availability and diversity are important drivers of global variation in fish trophic niches, with compression of the trophic niche occurring secondarily in more species-rich communities in response to current or past interspecific competition (Connell, 1980; MacArthur, 1970). Similar results have been observed for Maitland and Rahel (2023) at a smaller scale, where fish isospaces were determined by an interaction between environmental and species diversity. Here, isospace size declined sharply when basin richness exceeded 50 fish species. This may indicate some degree of functional redundancy in highly diverse communities (Loreau, 2004; Walker, 1992) or suggest that species-poor communities function differently from species-rich ones—being more shaped by stochastic processes, whereas richer communities are more deterministic. Metacommunity dynamics (e.g., mass effects, connectivity) may also influence how competition shapes trophic niche width (Bauer et al., 2021; Quévreux & Loreau, 2022). Further studies are needed to disentangle these mechanisms. However, we acknowledge that basin-scale species richness is not equivalent to species diversity at the spatial scale at which species interact, but data for local-scale fish diversity was unavailable for many of our datasets.

Terrestrial food resources subsidize many, if not most, freshwater communities (Lafage et al., 2019; Leal et al., 2023). During river flow pulses, inundated floodplains allow fish to access terrestrial invertebrates, fruits, seeds, leaves, and detritus (van der Sleen & Rams, 2023). Thus, fish trophic niches are likely to expand during high-flow pulses, particularly in tropical regions where wet/dry seasonality produces floods of large magnitude and duration annually. We used the CV of precipitation as a proxy for water level fluctuations in aquatic systems. This variable had a weak, positive association with population isospace size, supporting the EVH hypothesis. Several factors could have contributed to this weak association. First, we did not account for differences in productivity between aquatic and terrestrial ecosystems, which are known to influence the degree of terrestrial subsidies (Marczak et al., 2007). Second, our metric does not take into account landscape features influencing flood dynamics. In some cases, local precipitation may be an inadequate proxy for water fluctuations, for example when precipitation or snowmelt in upstream regions is the main driver of hydrology. Third, isotopic signatures may not capture seasonal variability in precipitation, as their turnover time is typically limited to a relatively short period—ranging from weeks to a few months (Vander Zanden et al., 2015). Finally, a lack of isotopic differentiation between aquatic and terrestrial food resources would hinder the ability to detect the effect of terrestrial subsidies on consumer isospace (Winemiller et al., 2022).

Fish population isospace differed among different types of ecosystems, which is consistent with the expectations of the ETH hypothesis. Fishes in large lakes had slightly smaller isospaces than those inhabiting small lakes, streams, and rivers. In large lakes, most fishes may be supported by pelagic food chains originating from phytoplankton (Maitland et al., 2024). By contrast, smaller water bodies, including small lakes and streams, have higher shoreline/area ratios, allowing for greater inputs of allochthonous material from surrounding terrestrial ecosystems, which in turn would increase food resource diversity for fishes (Larson et al., 2011). Rivers and streams are heterogeneous and dynamic ecosystems, and their food webs are supported by diverse energy sources differing in isotopic composition (Potapov et al., 2019; Riede et al., 2011; Roach, 2013). However, the relatively small differences in isospace among ecosystem types reflect substantial variability within each ecosystem that was not accounted for in the analysis. Among these, the interaction with marginal floodplain lakes and wetlands where methanogenic carbon assimilation may occur (e.g., Urbano et al., 2025) and changes in resource subsidies from headwaters to mouth (e.g., Soria-Barreto et al., 2020) may represent extreme isotopic variation at the base of food webs reflected to fish consumers.

### Isospace variation — Traits

Although trophic guild was the strongest predictor of fish isospace size, it did not support the predictions of either the TLH (Jennings et al., 2002; Layman et al., 2005; Scharf et al., 2000) or the OTH (Woodland et al., 2015) hypotheses. Instead, primary consumers tended to have larger isospaces than fishes at intermediate (invertivores, planktivores) and high trophic levels (carnivores and piscivores). Our finding agrees with a recent study of fish food webs in a subtropical river where species at lower trophic levels (TL < 3) had larger isospaces than predators (Zhang et al., 2024). The trophic niche of herbivorous and detritivorous/algivorous fishes has historically been underestimated, largely due to the difficulty of identifying minute and often fragmented or amorphous items in gut contents (Martinson et al., 2008). Stable isotope analysis provides an alternative approach to estimate trophic niches, even though considerations that specific diet compositions are unknown. Primary consumers should be selective foragers due to the high variability in the nutritional quality of plant tissues (Clauss et al., 2013; Martinson et al., 2008). However, large herbivorous and detritivorous fishes have high consumption rates, which may hinder the evolution of specialized diets (Arim et al., 2010; Sibly et al., 2012). Some studies have shown that herbivorous and detritivorous fish under certain conditions supplement their diet with invertebrates (e.g., Andrade et al., 2019; Nalley et al., 2021). Furthermore, herbivorous fishes may shift their diet from autochthonous to allochthonous plant material when highly nutritious forms of the latter become available during the flood season (Correa & Winemiller, 2013; Ou & Winemiller, 2016).

While a detritivore or herbivorous fish may appear to have a narrow dietary niche based on traditional gut content analyses—where "detritus" is often treated as a single food item—it may actually exhibit a broad isotopic niche. This is because detritus can originate from diverse sources and be subject to different isotopic fractionation processes. For instance, allochthonous detritus decomposed in anoxic, methanogenic habitats can have δ¹³C values below –35‰, while autotrophic detritus and algae may have δ¹³C values as high as –15‰. Additionally, in herbivorous fishes, stream current velocity can significantly influence algal δ¹³C signatures (Trudeau & Rasmussen, 2003), such that feeding in lentic versus lotic environments can result in markedly different isotopic values in consumer tissues. Therefore, when populations of herbivorous or detritivorous fish include individuals feeding in habitats with varying current velocities or oxygen conditions, their isotopic niche—or isospace—is expected to be wide.

The greater population isospace variability at the base of the food web may stem from the progressive loss of isotopic differentiation as material is transferred up the food chain. Consumers integrate the isotopic compositions of their assimilated resources, leading to intermediate δ¹³C values (Fry, 2007). Although δ¹³C is skewed toward the most important resource, values tend to average out at higher trophic levels. As a result, when plotted in δ¹³C– δ¹ N space, food webs often form a triangular pattern, with high δ¹³C variation at low δ¹ N (a proxy for trophic level) and a narrower range of intermediate δ¹³C values at high δ¹ N (Keppeler et al., 2021; Maitland et al., 2024). This pattern implies that top predators are unlikely to have an extremely depleted or enriched ¹³C signature. The effect may be particularly pronounced in size-structured food webs, as larger organisms typically have lower tissue turnover rates than smaller ones (Vander Zanden et al., 2015). Slower turnover allows the organism’s isotopic signatures to integrate assimilated material over extended periods and across different locations, further driving δ¹³C values toward intermediate levels (Keppeler et al., 2021). The extent to which trophic patterns may be biased by body size and differential rates of organic matter assimilation is unknown, especially since the triangular pattern is not evident in some food webs (e.g., Andrade et al., 2023). Future studies may consider using mixing models to refine the contribution of basal sources to the diet of consumers (Hoeinghaus & Zeug, 2008); however, these efforts are hindered by limited knowledge about trophic enrichment factors and tissue/elemental turnover rates (Bastos et al., 2017; Vander Zanden et al., 2015). Analysis of fatty acids (Haubert et al., 2011), compound-specific isotopes (Bowes & Thorp, 2015), and DNA barcoding (Wolfe et al., 2025) of ingested material could provide greater resolution of diet composition.

Our findings support the predictions of the BVH hypothesis, as intrapopulation variation in body size was associated with larger isospaces. Among the tested variables, body size variation emerged as the second strongest predictor. This relationship is attributed to ontogenetic niche shifts, in either feeding, habitat use, or both (Werner & Gilliam, 1984). In addition to phenotypic variation associated with ontogeny, the adult fish phenotypes have been shown to vary as a function of genetic and environmental factors (Ahti et al., 2020) as well as sexual dimorphism (Parker, 1992). Isospace variation could also be influenced by other factors, like territoriality, where larger individuals control spaces with the most profitable resources (Schirmer et al., 2019), resulting in significant between-individual niche variation (Bolnick et al., 2003).

The average body size of fish populations had a positive but weaker relationship with isospace size than variation in body size within populations. The positive trend is consistent with the BEH hypothesis, which suggests that trophic diversification occurs to meet higher energy demands and is facilitated by mouth-gape release (Arim et al., 2010; Rooney et al., 2008). However, the weak association may indicate that the hypothesis assumptions do not hold for some species (Keppeler & Winemiller, 2020a; Hanson et al., 2024) or simply stable isotopes fail to clearly detect the underlying mechanisms, particularly in distinguishing generalist from specialist individuals (Hette-Tronquart, 2019).

Contrary to the expectations of the SEH hypothesis, fish populations with truncated or round caudal fins exhibited broader isospaces than those with forked, emarginate, or lunate caudal fins. A possible explanation is that fish with lower vagility (truncated or round caudal fins) forage within smaller areas. If these individuals are more spatially segregated compared to more vagile species that seek preferred food resources over large areas, a broader isospace could result from the isoscape structure and/or individuals consuming different locally available resources. By contrast, fish with forked, emarginate, or lunate caudal fins are generally more vagile and may have isotopic signatures that integrate those of resources obtained over broad areas. Additionally, many species with forked caudal fins, such as South American piranhas (*Serrasalmus* spp.) and African tigerfish (*Hydrocynus vittatus*), are carnivorous and tend to have relatively small isospaces compared to fishes at lower trophic levels. Forked caudal fins are also common among schooling fishes (e.g., South American tetras), whose aggregative behavior tends to homogenize prey items among individuals and likely reduce population trophic niche width.

### Isospace variation — Scale

Although we set a minimum sample size for populations included in our analyses, we still observed moderate increases in isospace size with sample size, consistent with the SSH hypothesis. Additionally, isospace size tended to be larger when more habitats and years were surveyed, in agreement with the STH and TSH hypotheses, respectively. Notably, including additional habitats and years had a considerably greater effect than randomly sampling more individuals, emphasizing the need for standardized sampling protocols when comparing trophic niches of populations. Spatial and temporal variation in isotopic ratios and availability of resources as well as time lags in the assimilation of elements from ingested food items are factors affecting animal isotopic ratios (Fry, 2007; Layman et al., 2012). Our results underscore the importance of factoring out scale-dependent variables, even when adhering to the minimum sample sizes recommended in the literature (Eckrich et al., 2019; Jackson et al., 2011).

### Differences between predators and non-predators

Our findings suggest that the factors shaping isotopic niche space (isospace) differ between predatory and non-predatory fishes. Scale-dependent factors, for example, were nearly three times more important in explaining isospace variation among non-predators than among predators. Most of this variation was associated with habitat diversity, indicating that spatial differences in the isotopic signature of basal resources have a stronger influence on non-predatory species. This may reflect the greater spatial isolation of non-predators, which tend to be smaller in body size and rely on a narrower range of energy sources (Arim et al., 2010; Rooney et al., 2008). Another possible explanation is the higher variability typically found in the isotopic signatures of basal resources compared to those at higher trophic levels (Hoeinghaus & Zeug, 2008).

Non-predators also showed consistently different relationships with solar radiation and basin fish richness compared to predators. Basin richness was positively associated with isospace in non-predators, suggesting that interspecific competition may not be a major constraint on their trophic niche breadth. This may reflect the greater availability of energy at the base of the food web, where less energy is lost through trophic transfers (Thompson & Townsend, 2005). However, it is important to note that basin-level fish richness may not be a strong proxy for resource competition in non-predators, as their primary food sources—such as detritus and macrophytes—are also heavily exploited by other aquatic organisms, particularly invertebrates. In contrast, we observed lower isospace values in non-predators under high solar radiation. This could result from inhibited basal resource production or shifts in basal community composition under intense solar exposure, especially ultraviolet radiation. Such conditions may favor the proliferation of dominant producers like algal blooms (Paul, 2008) and free-floating macrophytes, which can drastically alter water chemistry and light penetration (Häder et al., 2007; Lind et al., 2022). Alternatively, increased productivity associated with solar radiation might lead consumers to concentrate their diets on the most abundant and nutritious resources, thereby narrowing their trophic niche (Peterson et al., 2017).

In contrast to predators, non-predatory fishes exhibited a decline in isospace with increasing body size and an increase with higher caudal fin aspect ratios. The smaller isospace observed in larger individuals may reflect the inverse relationship between body size and trophic position commonly found in non-predators (Burress et al., 2016; Keppeler et al., 2020). While smaller non-predatory species tend to incorporate more animal material into their diets—thus expanding their trophic niche—larger individuals are typically more specialized, often possessing long intestines adapted for digesting detritus and/or plant material (Burress et al., 2016; Keppeler & Winemiller, 2020a). The relationship between caudal fin aspect ratio and isospace differs between predators and non-predators, though the reasons remain unclear. This pattern may be related both to differences in the isotopic signatures of consumed resources— which are often more heterogeneous for herbivores (Hoeinghaus & Zeug, 2008)—and to the functional role of the caudal fin, which is more critical for predators that rely on speed and maneuverability to capture prey (Sambilay et al., 1990).

### Latitudinal gradient and ecoregion differences

Although fish isospaces tended to be smaller at higher latitudes, our results did not reveal a consistent latitudinal gradient. Isospaces were smaller in the Nearctic and Australasia. In Australasia, the sampled aquatic ecosystems were notably drier than those on other continents (Figure S1), and their fish populations were mostly supported by algae (Beesley et al., 2020; Bunn et al., 2013) and had relatively small isospaces. Most Nearctic fish populations were carnivorous species with relatively small isospaces. Higher proportions of predatory fishes at higher latitudes are a well-documented macroecological pattern (Moss, 2013), and greater incidence of herbivory and omnivory in (sub)tropical fishes (González-Bergonzoni et al., 2012) may explain the broader isospaces observed in the Neotropics and Indo-Malay ecoregions. The larger isospaces in Palearctic fish compared to Nearctic fish may be associated with the lower species richness and potential competition in the former as well as bias due to fewer studies included from that region.

### Caveats

A limitation of SIA and our isospace approach is that the isotopic space does not necessarily reflect the trophic niche with precision and accuracy. For instance, each individual of a population occupies a single point in isotopic space, whether or not that individual has a narrow or broad diet. Thus, if a population has low between-individual diet variation, the population isotopic space would be narrow even though individual diets are broad (Hette-Tronquart, 2019). In addition, δ^13^C, δ^15^N and other isotopic ratios of organisms can be influenced by factors not directly related to diet (Hette-Tronquart, 2019; Hoeinghaus & Zeug, 2008), including environmental variables (e.g., temperature, variation in inorganic elemental sources available for primary producers), anthropogenic impacts (e.g., sewage effluents, fossil fuel usage), lipid content, and consumer-associated traits (e.g., metabolism, growth rate) (Bastos et al., 2017; Davis et al., 2012; Hoffman et al., 2015; Maitland et al., 2021; Vander Zanden et al., 1997; Villamarín et al., 2018). Although it is impossible to account for all variables influencing stable isotope composition, we acknowledge these limitations and their potential effects on the global patterns observed.

### Conclusions

Analyzing population isospaces is complicated by various non-trophic factors influencing isotope ratios (Hette-Tronquart, 2019; Hoeinghaus & Zeug, 2008). Investigation of trophic niches at a global scale presents challenges, including data standardization and statistical approaches to control for potential confounding factors. Despite these limitations and challenges, our global analysis supports several ecological hypotheses, including productivity–diversity (PDH) and Body Size Variation-Trophic Expansion (BVH) hypotheses, and provides insights into new ones, such as Swimming Efficiency-Trophic Breadth hypothesis (SHE). Advancing our understanding of trophic niche breadth is important, not only because niche theory is central to ecology, but also because it informs predictions of species response to environmental variation, including changing conditions of the Anthropocene.

## ACKNOWLEDGMENTS

We thank the National Council for Scientific and Technological Development of Brazil (CNPq) for providing a postdoctoral scholarship (150773/2024-2) to FWK to support this research, as well as many other public (Fundação de Amparo e Desenvolvimento da Pesquisa, US National Science Foundation, US Environmental Protection Agency, Texas Parks and Wildlife Department, Carlos Chagas Filho Foundation for Research Support of the State of Rio de Janeiro, Brazilian Federal Agency for Support and Evaluation of Graduate Education, and The Scientific and Technological Research Council of Türkiye) and private institutions (Norte Energia, National Geographic Society, and Sabin Center for Environment and Sustainability) that funded field studies and stable isotope analyses. We also acknowledge the efforts of numerous volunteers, students, and technicians who collected and processed samples. The views expressed in this manuscript are those of the authors and do not necessarily represent the views or policies of the U.S. Environmental Protection Agency.

## DATA AVAILABILITY STATEMENT

The data will be deposited in Dryad following the acceptance of the paper for publication.

## AUTHOR CONTRIBUTIONS

Conceptualization: FWK, CGM, KOW; Funding acquisition: FWK, TG; Investigation (Data collection and processing): TG, CGM, AA, CCA, EB, TB, RE-S, CER, IG-B, DH, JH, OPJ, EJ, CL, EOL, BM, SSM, JAO, GP, YQ, AG, RKK, AT, PAT, TFT, EZ, KOW; Formal analysis: FWK; Visualization: FWK, TG; Writing - Original Draft: FWK, TG, CGM, KOW; Writing - Review & Editing: All authors.

